# Analysis of cortical cell polarity by imaging flow cytometry

**DOI:** 10.1101/2023.05.24.542074

**Authors:** Jesper Huitfeld Jespersen, Andras Harazin, Anja Bille Bohn, Anni Christensen, Esben Lorentzen, Anna Lorentzen

## Abstract

Metastasis is the main cause of cancer-related death and therapies specifically targeting metastasis are highly needed. Cortical cell polarity (CCP) is a pro-metastatic property of circulating tumor cells (CTCs) affecting their ability to exit blood vessels and form new metastases that constitutes a promising point of attack to prevent metastasis. However, conventional fluorescence microscopy on single cells and manual quantification of CCP are time-consuming and unsuitable for screening of regulators. In this study, we developed an imaging flow cytometry (IFC)-based method for high-throughput screening of factors affecting CCP in melanoma cells. The artificial intelligence (AI)-supported analysis method we developed is highly reproducible, accurate, and orders of magnitude faster than manual quantification. Additionally, this method is flexible and can be adapted to include additional cellular parameters. In a small-scale pilot experiment using polarity-, cytoskeleton-or membrane-affecting drugs, we demonstrate that our workflow provides a straightforward and efficient approach for screening factors affecting CCP in cells in suspension and provide insights into the specific function of these drugs in this cellular system. The method and workflow presented here will facilitate large-scale studies to reveal novel cell-intrinsic as well as systemic factors controlling CCP during metastasis.

## Introduction

While metastasis is the major cause for cancer-related deaths, cancer treatments still largely target primary tumors and not metastatic spreading (Ganesh & Massagué, 2021). One reason for this discrepancy is that cancer treatment success has been evaluated based on primary tumor shrinkage and not inhibition of metastasis. The Food and Drug Administration has recently included metastasis-free survival as an endpoint for clinical trials, offering new prospects to advance development of anti-metastatic treatments for cancer patients (Alečković et al., 2019; Fernandes et al., 2019; Ganesh & Massagué, 2021).

During distant-organ metastasis, cells can pass through a liquid phase in lymph or blood as circulating tumor cells (CTCs) (Massagué & Obenauf, 2016; Ring et al., 2023). To form secondary tumors, CTCs need to survive the harsh conditions in blood circulation and extravasate from the blood stream through the vessel wall (Massagué & Obenauf, 2016; Ring et al., 2023). In circulation, CTCs are exposed to potential drugs and can be isolated and monitored in patients by minimally invasive blood sampling (liquid biopsies). Circulation and extravasation therefore constitute particularly promising points of attack to interrupt metastasis.

Extravasation is an active process of cell attachment, adhesion and migration that requires polarization and directionality (Estecha et al., 2009; Strilic & Offermanns, 2017). We have previously described a type of cortical cell polarity (CCP) in CTCs that constitutes a basic polarity module restricted to the plasma membrane (PM) and the sub-membrane actin cytoskeleton (Lorentzen et al., 2018). This pole is characterized by folding of the PM and local accumulation of filamentous actin and the linker protein ezrin. CCP promotes attachment and adhesion of CTCs *in vitro* and metastatic seeding in animal models. CCP thus constitutes a potential target to reduce metastasis by interfering with the ability of CTCs to extravasate (Heikenwalder & Lorentzen, 2019; Ring et al., 2023).

Methodically, CCP has been assessed by widefield imaging and qualitative inspection or tedious manual quantification of polarized cells (Lorentzen et al., 2018). As this microscopy-based method is impractical for medium- or large-scale screens for regulators of CCP, we here developed methodology with comparably sensitive but faster readout combined with robust automated quantification.

Imaging flow cytometry (IFC) combines the spatial resolution of microscopy with the speed and quantitative multiparameter analysis of flow cytometry (Rees et al., 2022). On the ImageStreamxMk II (Cytek Biosciences), used in this study, each event is identified based on an image captured by a CCD camera, on a pixel-by-pixel basis in up to 12 channels. The technique provides fluorescent and morphological parameters of up to 5000 cells/second and is therefore a powerful method for automated, high throughput analysis of cellular and subcellular features of cells in suspension.

We established an IFC-based workflow as high-throughput method to measure CCP based on ezrin-GFP localization and present a reproducible artificial Intelligence-(AI) supported image analysis algorithm to quantify cortical polarization. As proof-of-principle, we conducted a small-scale pilot screen of compounds with potential effects on CCP. Our results demonstrate robust and reliable measurement of CCP in melanoma cells and show that this workflow can be adapted to include measurements of additional cellular features.

## Materials & Methods

### Cell culture

Generation and use of SkMel2 melanoma cells (ATCC: HTB68) stably expressing ezrin-GFP (SkMel2-ezrin-GFP), was described previously (Lorentzen et al., 2018). Cells were grown at 37°C under 5% CO_2_ in high glucose Dulbecco’s modified Eagle’s medium with Glutamax (Thermo Fisher Scientific) supplemented with 10% fetal bovine serum (FBS, Merck), 100U/ml Penicillin and 100μg/ml Streptomycin (Thermo Fisher Scientific) and 1mg/ml active G418 (Merck). For maintenance, cells were detached with 0.5g/l trypsin and 0.2g/l ethylenediaminetetraacetic acid solution (EDTA) (Merck), for experiments, with 0.2g/l EDTA (Merck). Phosphate buffered saline (PBS) (Merck) was used for washes. Cell numbers and viability were measured using a NuceloCounter NC-200 (Chemometek).

### Chemicals and drugs

Chemicals were dissolved in dimethyl sulfoxide (DMSO) (Merck) or 96% ethanol (VWR) according to the manufacturers recommendations, further diluted in cell culture medium and used at the following concentrations: Forchlorfenuron (Merck) at 100μM, cholesterol sulfate (Merck) at 35μM, NSC23766 (Tocris Bioscience) at 10μM, Dasatinib (Merck) at 0.5μM, Ml141 (Tocris Bioscience) at 1μM. Control experiments contained 1:500 DMSO or 1:500 ethanol in cell culture medium. 16% formaldehyde solution (Thermo Fisher Scientific) was added to suspension cultures to an end concentration of 4%. 4′,6-diamidino-2-phenylindole (DAPI, Thermo Fisher Scientific) was used at 0.2μM.

### Polarity Assays

SkMel2-ezrin-GFP cells grown in 10cm diameter dishes (VWR) to 80-90% confluency were treated with the indicated agents in medium for 24 hours, washed, detached with EDTA for 15 minutes at 37°C, resuspended in medium and incubated in suspension for the indicated time (15 minutes, 1 hour or 3 hours) at room temperature for the training data sets or resuspended in medium containing the indicated reagent for 1 hour at room temperature for IFC experiments. After incubation, cells were fixed in 4 % formaldehyde for at least 1 hour, washed with PBS, resuspended in PBS containing 0.2μM DAPI and maintained at 4°C until measurement by IFC or mounted in Mowiol (Merck) on a microscopy slide.

### Fluorescence microscopy

Manual polarity assays were performed on an IX83 widefield microscope (Olympus/Evident, Tokyo, Japan) with a halogen lamp using 10x (Fig. 1B) or 20x (Fig. 3C) objective and standard FITC filters. Exposure time was set to 200 ms, gain to 3 in CellSens Standard software (Olympus/Evident). Manual quantification was done on 3 (Fig. 3C) or 4 (Fig. 1B) images containing at least 200 cells per replicate. Confocal microscopy was performed on an LSM710 microscope (Zeiss, Oberkochen, Germany) with multialkali PMTs using a 63x 1.4NA oil immersion objective. DAPI was excited using a 405 Diode laser, GFP using the 488-laser line of an Argon laser. Images were recorded using ZEN black software (Zeiss) and processed in Fiji (Schindelin et al., 2012).

**Figure 1:**
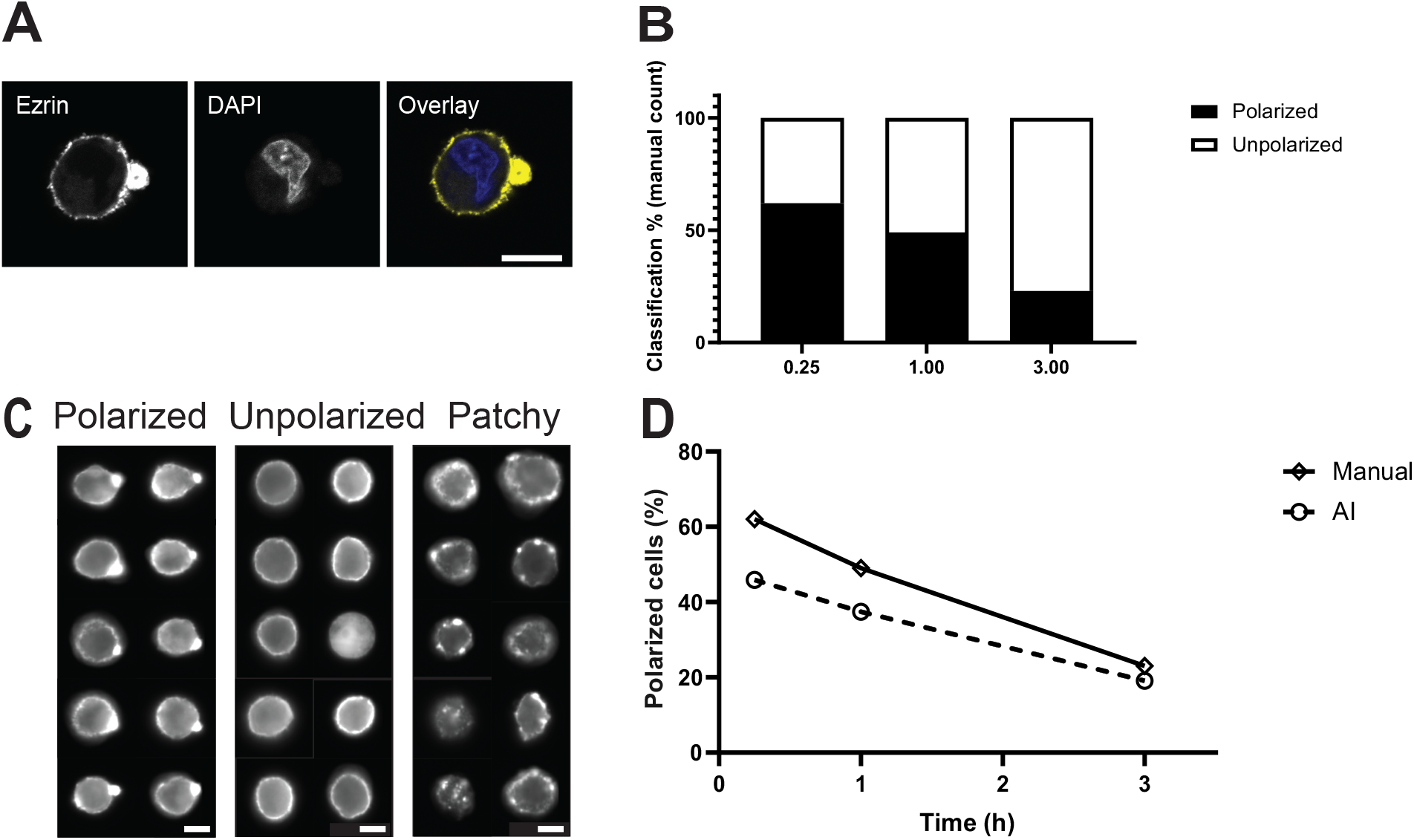
Method development. **A**. Localisation of ezrin-GFP in an SkMel2 cell displaying CCP imaged by confocal microscopy. Scale bar = 10μm. **B**. Quantification of the fraction of SkMel2 cells displaying CCP (black) as measured by polarized ezrin-GFP distribution using widefield microscopy and manual quantification. For the training data set, cells were maintained in suspension for 15 minutes (0.25), 1 hour (1.00) or 3 hours (3.00). Single measurements are shown. **C**. Images obtained from SkMel2-ezrin-GFP cells by IFC using 60x magnification. Cells are manually classified into polarized (left), unpolarized (center) or patchy (right). **D**. Comparison of the manual quantification (solid line, displaying the same data sets as in **(B)**) and AmnisAI-supported (dashed line) classification of the fractions of polarized cells.

**Figure 2:**
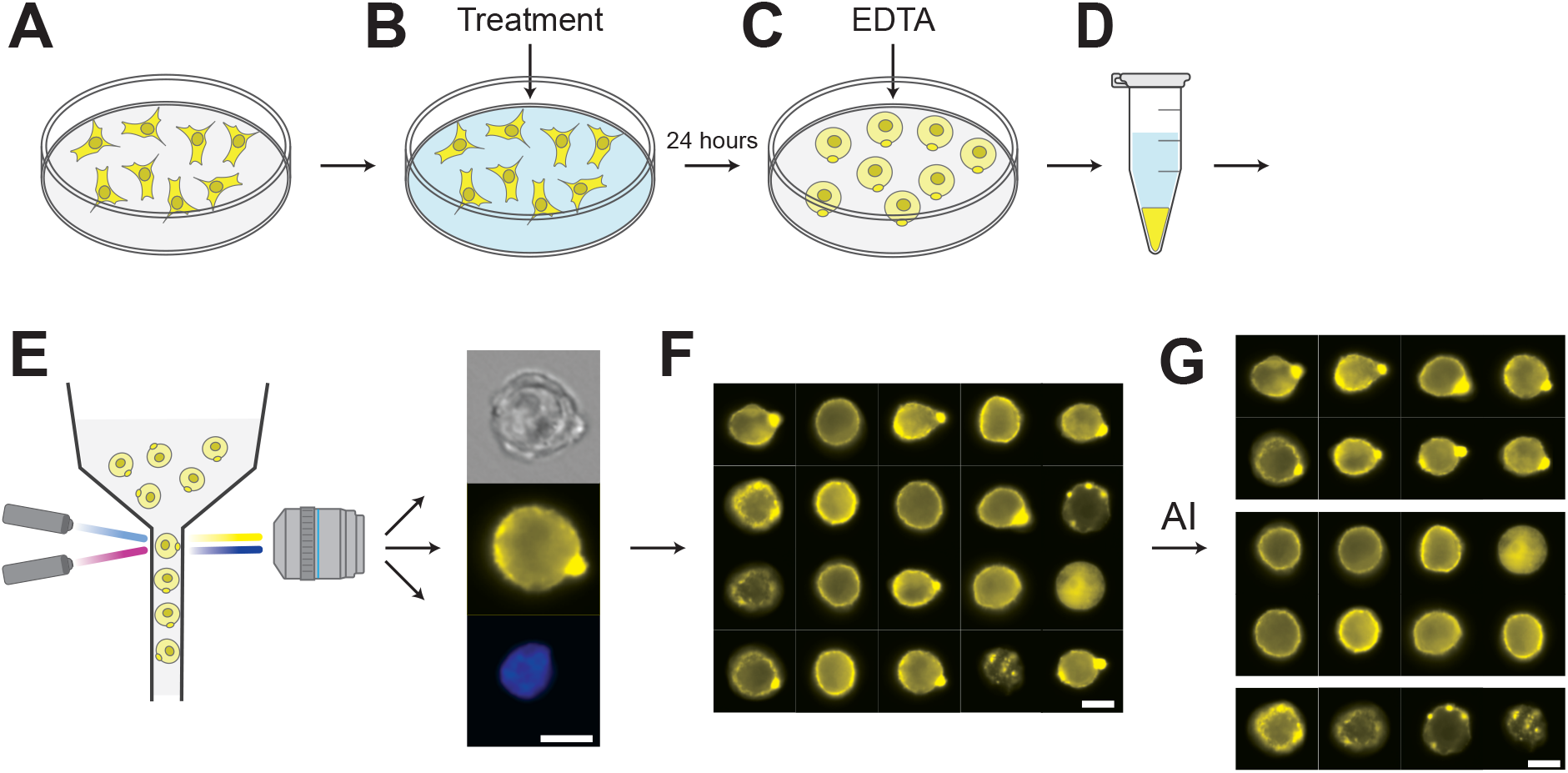
Workflow. Illustration of the IFC- and AI-based method to determine the fraction of cortically polarized cells. **A-D** shows polarity assays and **E-G** IFC and image analysis. SkMel2-ezrin-GFP cells cells are grown in adhesion cultures **(A)**, treated with potential polarity-modulating agents **(B)**, detached **(C)** and incubated and fixed in suspension **(D)**. Images of 40000 cells are recorded by IFC **(E, F)**, gated in IDEAS and classified into polarized, unpolarized and patchy subpopulations **(G)** by AmnisAI.

**Figure 3:**
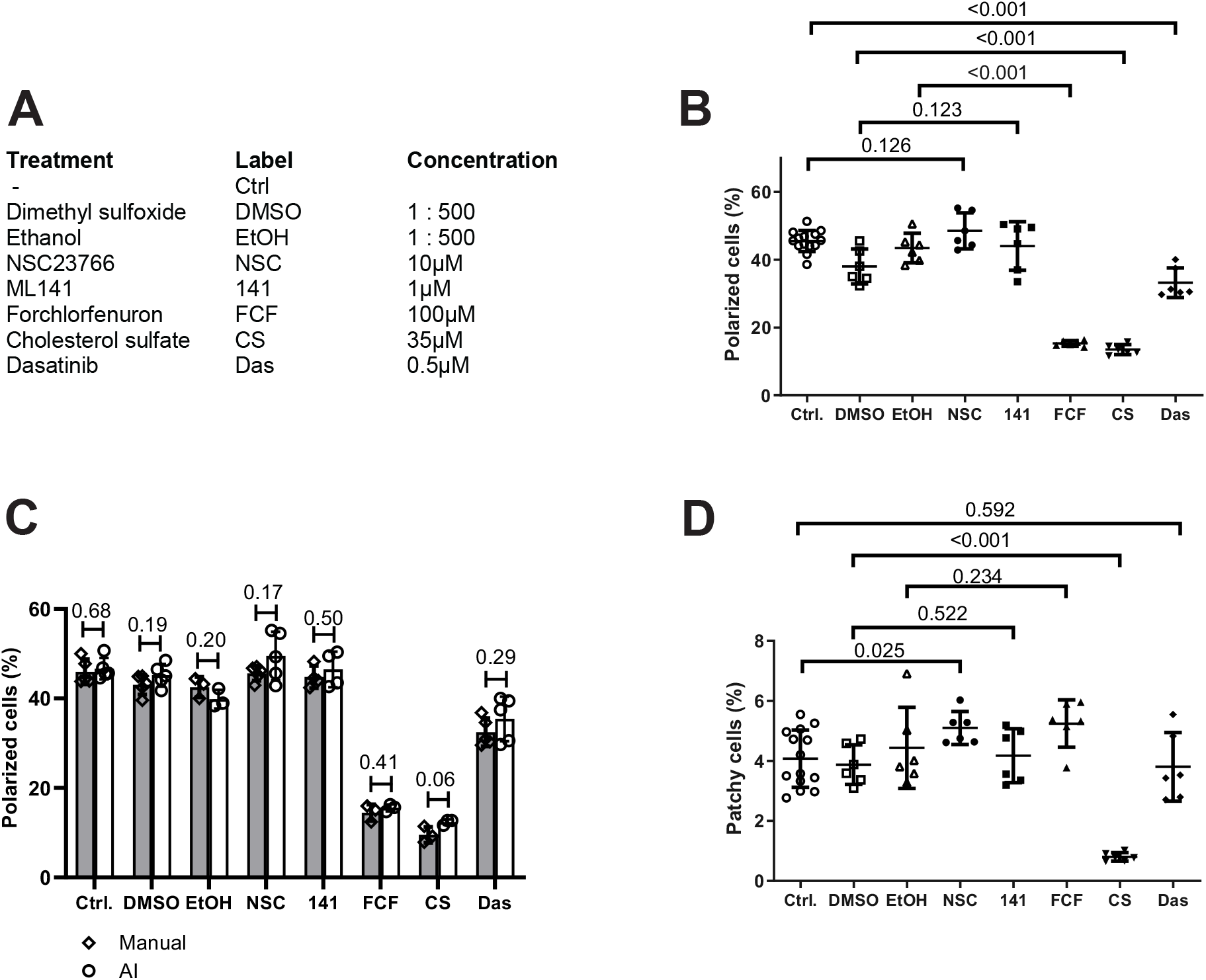
Proof-of-concept experiments. Using the workflow in Figure 2, SkMel2-ezrin-GFP cells were subjected to the indicated treatments and the effect on CCP was determined by IFC. **A**. Treatments and concentrations used for this screen. **B**. Fraction of cortically polarized cells as assessed by IFC method after indicated treatments. Individual measurements with mean ± SD, n=6, p-values from unpaired t-tests. **C**. Three randomly selected experiments from data shown in **(B)** were quantified by widefield microscopy and manual count (grey bars) and compared to AmnisAI classification (white bars). Individual measurements and bar diagrams ± SD, n=3, p-values from unpaired t-tests. **D**. Quantification of the fraction of SkMel2 cells displaying patchy ezrin-GFP distribution classified by AmnisAI in the experiments shown in **(B)**. Individual measurements with mean ± SD, n=6, p-values from unpaired t-tests.

### Imaging flow cytometry

Imaging flow cytometry was performed on an Amnis ImageStream®XMkII (Cytek Biosciences, Fremont, CA, USA) at 60x magnification at slow speed (for highest quality). DAPI was excited by 405nm laser (70 mW) and detected in channel 7, GFP was excited by 488nm laser at 70-90 mW and detected in channel 2. 488 nm laser power was adjusted for each experiment that untreated control samples contained ≈4% oversaturated cells. Brightfield images were collected in channel 1 and 9. For each sample, 40000 cells were imaged. A compensation matrix was created in AmnisIDEAS software using images of 10000 untransfected DAPI-stained SkMel2 cells and 10000 SkMel2-ezrin-GFP cells and applied to all samples. Images were gated in IDEAS to exclude out-of-focus cells, doublets, double-nucleated cells, oversaturated cells in the GFP channel and GFP-negative cells. Apoptotic cells were identified as described by (Henery et al., 2008) and excluded. Approximately 50 % of cells were included for further analysis after gating and loaded in AmnisAI software for training of AI models or classification.

### AI model training

Initial base truth populations were selected in IDEAS. Only channel 2 GFP-images were included for training. AmnisAI was first trained based on the separation made in IDEAS and the resulting algorithm used to classify the cells. Cells that were by visual inspection deemed incorrectly classified, were manually moved to the appropriate truth population and the AI was retrained. This processes was repeated until no incorrectly classified cells were detected by eye. Performance of the algorithms was tested by calculating Precision, Recall and F1 values as:

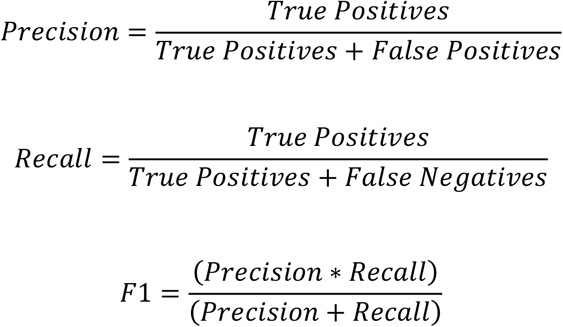

The AmnisAI model (.aimdl) files are available on request from the corresponding authors.

### Software and statistical analysis

AmnisIDEAS 6.3 (Cytek Biosciences, Fremont, CA, USA) was used for visualization, compensation and gating of IFC data. Amnis^®^AI 2.0.7 (Cytek Biosciences, Fremont, CA, USA) was used for training and classification of polarized, unpolarized and patchy cells and develop the analysis algorithm. CellSens (Olympus/Evident, Tokyo, Japan) and ZEN black (Zeiss, Oberkochen, Germany) were used for acquisition and Fiji (Schindelin et al., 2012) for processing of fluorescence microscopy images. Prism 9.5.0 (GraphPad, Dotmatics, Boston, MA, USA) was used for visualisation and statistical analyses of data. P-values are based on two-tailed non-paired t-tests with assumed Gaussian distribution.

## Results

### Establishment of an IFC-based method for measuring CCP

CCP is defined as a type of polarity adopted by cells in liquid phase without directional stimulus and is detected by formation of an ezrin-enriched pole (Lorentzen et al., 2018). We thus used ezrin-GFP localization as a marker of CCP in SkMel2 human melanoma cells in suspension (Fig. 1A). To establish a new IFC-based method, a template data set was generated using a published microscopy-based method (Lorentzen et al., 2018). Using the observation that CCP decreases over time, polarity assays were performed with SkMel2-ezrin-GFP cells maintained in suspension for 15 minutes, 1 hour or 3 hours after detachment. The fractions of polarized cells, as manually quantified using the microscopy-based method were 62%, 49% and 23% respectively (Fig. 1B). The same cell batches were subjected to IFC, recording at least 40000 cells for each condition. IFC provided high-quality images with clearly discernible polarized or unpolarized ezrin-GFP localization (Fig. 1C). Pre-processing steps including compensation and gating, were performed to remove out-of-focus images, cell duplets, apoptotic cells, non-GFP expressing cells and overexposed cells from analyses (Supplementary Fig. 1).

Attempts to classify polarized and unpolarized cells by manual adjustment of parameters and a machine learning function available in IDEAS were not successful, as they were not reflecting the results obtained by the manual quantification method, which is the standard that initially defined CCP (Supplementary Methods and Supplementary Fig. S2). We therefore employed AmnisAI software that allowed for manual corrections and training of the AI model to optimize classification. Generation of truth populations and AI training were performed on pooled data from the three time points shown in Figure 1B. Three truth populations (Fig. 1C) were generated by manually classifying polarized (1912 cells with one or two poles), unpolarized (1817 cells with unpolarized ezrin at the PM and unpolarized ezrin in the cytoplasm) and a third class termed ‘patchy’ (445 cells containing several small ezrin spots of different intensity distributed throughout the PM). The three final classes were established from the initial truth populations supported by several training rounds with manual corrections until no wrongly classified cells were observed. When this AI-supported classification was applied to our test data set (15 minutes, 1 hour and 3 hours), the resulting fraction of polarized cells reflected the manual quantification of the microscopy-based method resulting in 45.9 %, 37.4 % and 19.1 % polarized cells as compared to the respective 62 %, 49 % and 23 % polarized cells quantified by initial manual count (Fig. 1D). Precision values of 97.5%, 94.9% and 83.3% and Recall values of 92.1%, 97.7% and 93.3% were calculated for polarized, unpolarized and patchy classification, respectively, confirming that the AI-generated algorithm can classify our data with high confidence (Supplementary Fig. S3). In summary, a workflow has been established that allows for straightforward measurement and analysis of CCP (Fig. 2).

### Experimental evaluation of the quantification algorithm

To evaluate if the quantification algorithm can be applied to biological studies of polarity regulators, we performed a small-scale pilot experiment using a collection of agents that potentially affect polarity, the PM or the sub-membrane cytoskeleton (Fig. 3A). For our experiments, we selected two Rho-family GTPase regulators, the Rac1 inhibitor NSC23766 (NSC) (Gao et al., 2004) and the Cdc42 inhibitor ML141 (Hong et al., 2013), the septin inhibitor forchlorfenuron (FCF, 1-(2-chloro-4-pyridyl)-3-phenylurea) (Hu et al., 2008), the membrane component and signaling regulator cholesterol sulphate (CS) and the multitarget tyrosine kinase inhibitor dasatinib (Lindauer & Hochhaus, 2018). To this end, SkMel2-ezrin-GFP cells were pre-treated with NSC, ML141, FCF, CS, dasatinib or the respective solvent controls for 24 hours, subjected to polarity assays, imaged by IFC and classified using our AmnisAI algorithm (Fig. 2). In addition to polarization measurements, analysis of the DAPI channel in IFC allowed for identification of apoptotic cells (Henery et al., 2008). At the concentrations used in our experiments, none of the drugs or controls enhanced the fraction of apoptotic cells above the cut-off set at 1.0 % (Supplementary Fig. S4). We did not observe a decisive effect of NSC or ML141 on the fraction of polarized cells (Fig. 3B). Both FCF and CS strongly decreased the fraction of polarized cells from 45.6 ± 3.2 % in the untreated control to an average of 15.5 ± 0.8 % and 13.5 ± 1.5 %, respectively. Dasatinib decreased the fraction of polarized cells to an average of 33.2 ± 4.4 %. In summary, this pilot screen shows that the IFC-based measurement and our AmnisAI-supported classification model are well suited to measure differences in the fractions of cortically polarized cells under different treatment conditions. To compare the IFC-based method to the microscopy-based method under treatment conditions and to assess the sensitivity of the two methods, three random samples used for the IFC experiments were additionally imaged by microscopy and counted manually (Fig. 3C). Comparison of the two methods showed little difference in the fractions of polarized cells (Fig. 3C), demonstrating that the IFC-based method and automated quantification is comparable in capturing changes in CCP to the microscopy-based method with manual quantification.

In addition to classifying polarized and unpolarized cells, the automated quantification includes a patchy class, which is not accounted for in the manual count. The fraction of patchy cells remained relatively low at 4.1 ± 1.0 % in the untreated control and was slightly increased to 5.1 ± 0.5 upon NSC treatment (Fig. 3D). In CS-treated cells, the fraction of patchy cells was reproducibly strongly reduced to 0.8 ± 0.1 %, indicating that patchy cells constitute not only a fraction of non-classifiable cells but a biological cellular phenotype.

### Introduction of PM-localized and cytoplasmic ezrin classes

Ezrin activity is regulated by phosphorylation and membrane binding (Pelaseyed et al., 2017) and active ezrin is both phosphorylated and membrane-bound. We set out to test whether it is possible to quantify the amount of membrane-bound ezrin and get a measure of ezrin activity in the images previously used to measure CCP. To this end, two new classes, PM-localized and cytoplasmic, were introduced and analyzed from our previously generated data. The new classes were generated from two new truth populations made from the previously used 15 minutes, 1 hour and 3 hours merged data, membrane-localized (1576 cells with ezrin localized at the PM) and cytoplasmic (1994 cells with cytoplasmic ezrin), (Fig. 4A). Both truth populations included polarized as well as unpolarized cells. A new AI model based on these truth populations was trained to sufficient confidence in classification (Suppl.Fig. S5) and applied to our full datasets (Fig.4B). FCF treatment increased the fraction of cells with membrane-bound ezrin, suggesting that FCF treatment enhances activity of ezrin while dasatinib treatment reduced the fraction of cells with membrane-bound ezrin indicating that dasatinib reduces activity of ezrin. Since the pole is by definition localized at the PM, we tested if polarization affects the classification of PM-localized and cytoplasmic ezrin. To this end, polarized and unpolarized classes were analysed separately for cytoplasmic or PM-localized ezrin (Fig.4C and 4D). As expected, PM-localized ezrin was in general higher but differences were less pronounced in the polarized fraction. While the effect of FCF treatment enhancing membrane-bound ezrin was confirmed in both analyses, the reducing effect observed for dasatinib was not confirmed. As the pole was initially defined by localization at the cortex/PM, the biological conclusions that can be drawn from these findings are limited and this analysis should be considered a proof-of-principle experiment to test the flexibility of the method. However, our analyses indicate that inhibition of septins by FCF enhances membrane-association of ezrin, reflecting its activation state. This second level of analysis demonstrates how simple new classifications and sub-classifications can be added and implemented in the workflow.

**Figure 4:**
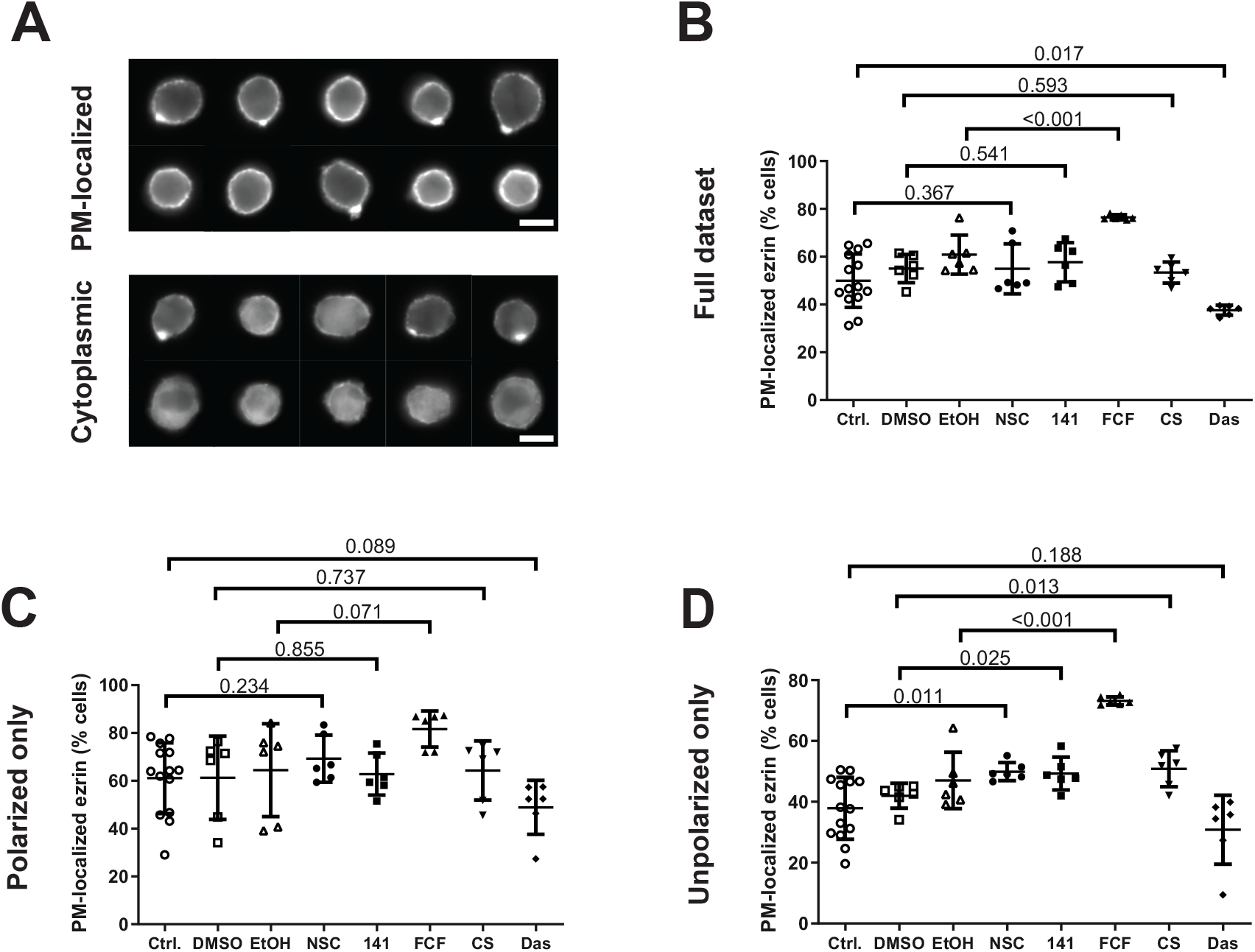
Subclassification of PM-localisation. The data displayed in Figure 3 were further analysed for PM- or cytoplasmic localisation of ezrin-GFP. **A**. Images obtained from SkMel2-ezrin-GFP cells by IFC using 60x magnification. After gating (Suppl. Fig. 1), cells were handpicked in IDEAS as PM-localised (left) or cytoplasmic (right). **B-D**. Fractions of cells classified as displaying PM-localised ezrin-GFP. Individual measurements with mean ± SD, n=6, p-values from unpaired t-tests. Subclassifications were performed on the full data set including polarized, unpolarized and patchy cells **(B)**, on cells previously classified as polarized **(C)** and on cells previously classified as unpolarized **(D)**.

Altogether, our results show that IFC-based measurement with automated analysis is comparably sensitive to microscopy-based measurement with manual counting in determining fractions of cortically polarized cells. In addition to being considerably faster and -once established-unaffected by human bias, the IFC-based analysis allows for straightforward introduction of additional measurements (apoptosis) or classes (patchy, cytoplasmic vs. PM).

## Discussion

We have established a straightforward workflow to quantify CCP. We demonstrated that this method is equally sensitive as a previously used microscopy-based method when used for biological screens of polarity regulators in tumor cells while being unbiased and automated. Importantly, this method is several orders of magnitude faster, which makes it applicable for medium-to large scale screens allowing for unbiased search for novel CCP regulators instead of limited directed approaches testing low numbers of suspected targets.

In this work, IFC and AI-supported image analyses were used to measure how drug treatments affect the percentage of cortically polarized cells. In addition, using a previously published method (Henery et al., 2008), the percentage of apoptotic cells was determined in each experiment without requiring additional measurements. In the cause of optimizing the classification, an additional phenotype termed ‘patchy’ was discovered. A similar distribution of multiple F-actin polymerization spots was described at the cortex of T cells during immune synapse formation upon microtubule depolymerization (Pineau et al., 2022). The patchy fraction was affected by treatments with CS and NSC, indicating that it constitutes a *bona fide* biological phenotype to be further investigated. We demonstrated that additional parameters (such as PM or cytoplasmic localization) can be straightforwardly implemented by AI training and analysed in the whole population or in previously generated subpopulations, to extract additional information from a single dataset.

For method evaluation, a small-scale pilot screen with regulators of actin or septin cytoskeleton, PM or cell signaling, was performed. Rho family GTPase regulators NSC (Gao et al., 2004) and ML141 (Hong et al., 2013) had no decisive effect on CCP, which allows for different interpretations. Firstly, Rac1/Cdc42 may not regulate CCP, which agrees with findings that the RhoA-specific guaninenucleotide exchange factor (GEF) GEF-H1 regulates CCP in early lymphocyte polarization during immune synapse formation (Pineau et al., 2022). Secondly, Rac/Cdc42 may need to act locally at the pole or outside the pole and global inhibition therefore shows no effect. This is supported by the finding that the patchy fraction of cells increased upon NSC treatment, showing that global Rac1 inhibition affects overall but not local cortical distribution of ezrin and possibly actin. Thirdly, NSC only inhibits interaction of Rac1 with the GEFs Trio and Tiam1 (Gao et al., 2004) and other Rac GEFs and GAPs might be involved in CCP. The specific function of actin regulators in CCP needs to be tested using systems-directed approaches taking the entire GEF/GAP/GDI regulatory network as well as subcellular local activities of Rho GTPases into account. The method presented here can facilitate screening of drug libraries targeting actin-regulating networks.

A strong reduction of polarized cells was observed upon FCF treatment, indicating a contribution of septins to CCP regulation. Septins are organisers of the cell cortex and regulators of the actin cytoskeleton and are directly associated with PM structures (Bridges & Gladfelter, 2015). A recent study revealed that membrane curvature at sites of PM blebbing induces cortical nucleation of septin and suggested that repeated blebbing may lead to stabilisation and formation of higher-order structures (Weems et al., 2023). According to this model, membrane blebbing and nucleation of septins could actively initiate or stabilize cortical polarization. Interestingly, septins are also involved in loss of polarity in aging hematopoietic stem cells (Kandi et al., 2021), indicating similarities between these two types of polarization.

Treatment with the cholesterol analogue CS reduced CCP, indicating regulation by either cholesterol-regulated signaling and/or the membrane lipid composition. Cholesterol affects mechanical cell properties via the actin cytoskeleton (Sun et al., 2007) and acts as naturally occurring membrane component but is also an inhibitor of the Rac-GEF DOCK2 (Sakurai et al., 2018). In addition, CS could affect polarization driven by PM lipid content or distribution, a hypothesis supported by the observation that the integrity of cholesterol-dependent PM microdomains is essential for the organisation of molecular cell polarity in hematopoietic stem cells (Görgens et al., 2012).

Dasatinib treatment reduced the fraction of polarized cells, implying tyrosine kinase signaling pathways in the regulation of CCP. Dasatinib is an anti-leukaemia drug that was developed as dual Src-Abl1 kinase inhibitor but targets a wide range of tyrosine kinases (Lindauer & Hochhaus, 2018). The specific tyrosine kinase pathway responsible for the observed effect can therefore not be distinguished. However, inhibition of cortical polarization by dasatinib may contribute to its anti-tumor effects.

In summary, we developed a simple protocol for IFC-based measurement and AI-based quantification of CCP directly applicable to screening for polarity regulators in tumor cell lines. Here, we only used two fluorescence channels (ezrin-GFP and DAPI) to extract multiple layers of data using advanced image analysis. The strength of IFC, however, lies in the possible acquisition of multiple fluorescence channels simultaneously, with each additional channel adding potential for multiple layers of image analysis. Our protocols constitute a basis to build on for further analyses.

The workflow presented here can be directly used to screen for regulators of CCP in cells in suspension. In addition to assessing CCP in cancer cell lines, this method can be adapted to measure CCP and other biologically relevant phenotypes of circulating tumor cells or immune cells, for monitoring in patients as well as basic scientific research. AI-supported analysis allows for fast and easy adaptation and limitless possibilities to combine polarity measurements with additional phenotypes. Altogether, this method opens the possibility of screening large libraries of molecules with unknown functions to identify new regulators and reveal possible pharmacologic inhibitors of CCP and help design potential anti-metastatic treatments.

## Acknowledgements

This work was funded partially by grants from the Novo Nordisk Foundation (grant number NNF20OC0064485) and Riisfort Fonden (E.L.) as well as Harboefonden, Brødrene Hartmanns Fond, Læge Sofus Carl Emil Friis og hustru Olga Doris Friis’ Legat and Tømrermester Jørgen Holm og Hustru Elisa f. Hansens Mindelegat (A.L.). We thank the FACS core facility and the Bioimaging Core facility at the Biomedicine Department at Aarhus University for providing access to instrumentation, training and support.

