## Supplementary material for "Analysis of cortical cell polarity by imaging flow cytometry": Suppl Figures and Methods

#### Supplementary Information

Supplementary Figures S1 - S5

Supplementary Methods

### Supplementary Figure S1

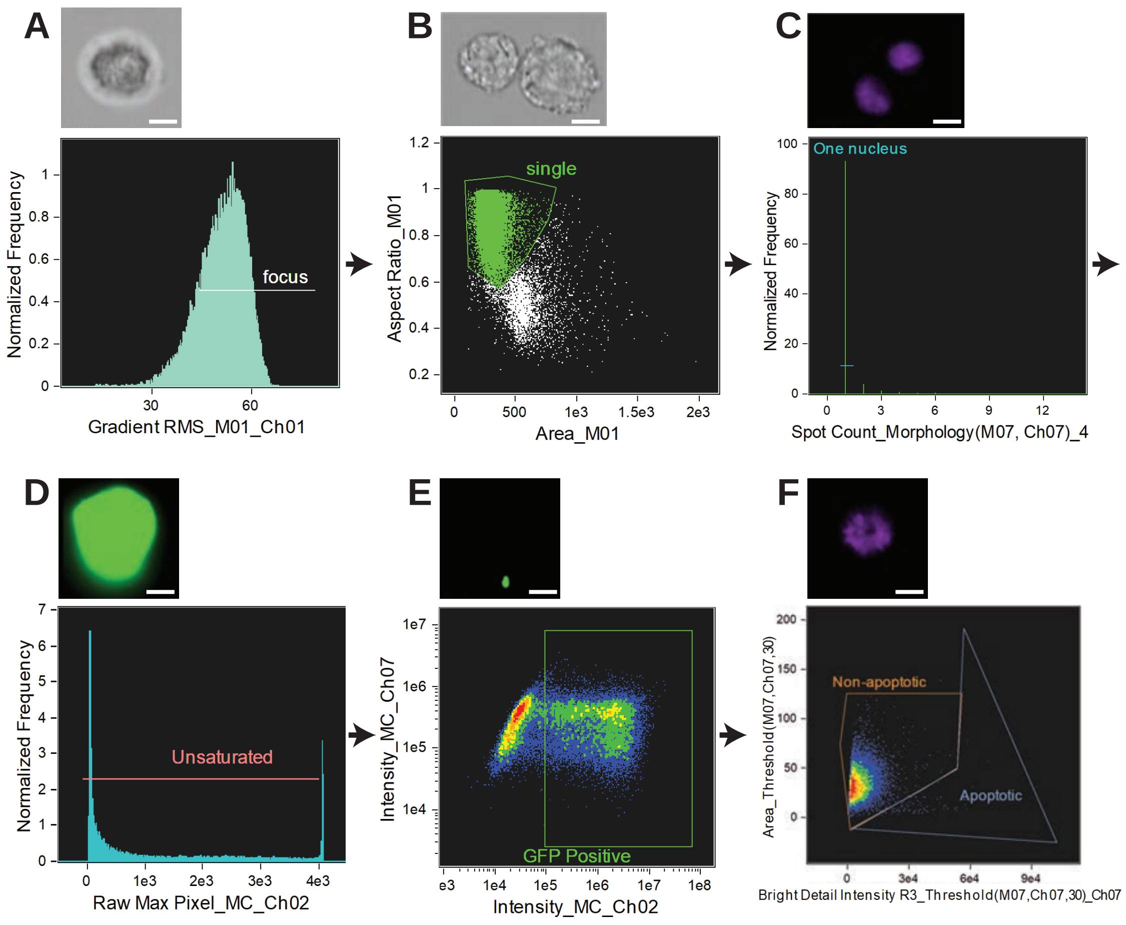


**Supplementary Figure S1: Gating strategy overview.**
The depicted gating strategy was implemented to identify cells to be included in the AI-based analysis before identifying truth populations. An example of the excluded cell type is shown on top of each plot (scalebar=7 µm). The gating strategy is described in detail as Supporting information in Supplementary Methods.
**A.** To select cells in focus, unfocused cell images were excluded from the analysis (Gradient RMS in the brightfield channel 1 < 42).
**B.** To select single cells, doublets were excluded from further analysis in a cell area versus aspect ratio plot.
**C.** Remaining doublets or double nucleated cells were excluded by excluding event with more than one nucleus.
**D.** Cells with oversaturated pixels in channel 2 (GFP) (Raw Max Pixel > 4094) were excluded.
**E.** GFP-positive cells were selected by gating on positive events in channel 2.
**F.** Apoptotic cells were removed from the analysis as described in Henery et al., 2008.

### Supplementary Figure S2

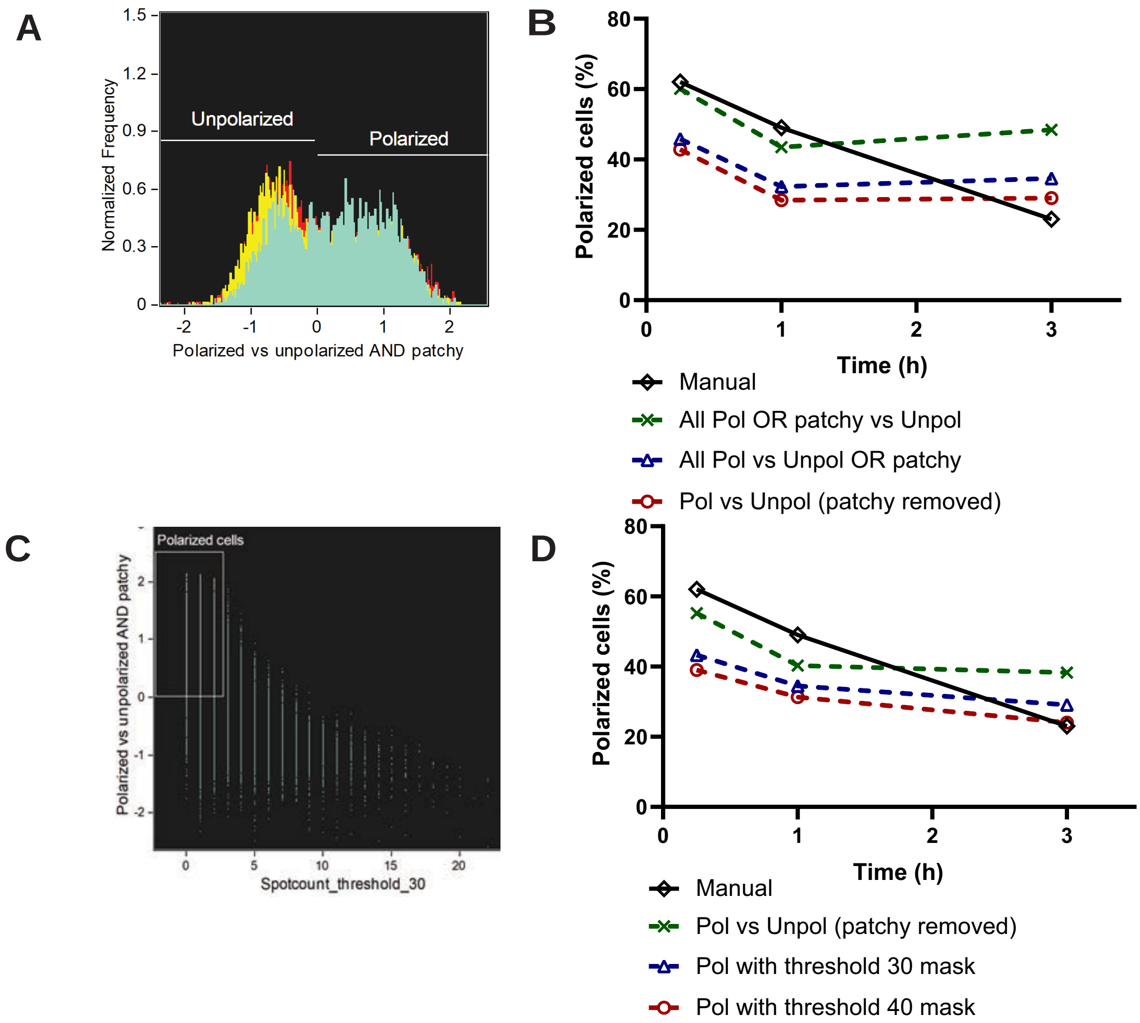


**Supplementary Figure S2: Machine learning and manual classification.**

**A**. Example of machine learning-based classification separating polarized cells from unpolarized and patchy cells in IDEAS software.

**B.** Fraction of polarized cells in the 15 min, 1 hour and 3 hour populations after classification using three different machine learning-generated classifiers of different combinations of polarized, unpolarized or patchy cells (dashed lines). 'Manual' indicates manual quantification after microscopy (data from Fig. 1B) that was used as a template. None of the machine learning-generated classifiers were able to reflect the decrease from the 1 hour to the 3 hours time point.

**C**. Example of a combination of a machine learning classifier with a user-defined IDEAS feature. This example shows a Pol vs Unpol OR patchy machine learning classifier on the y-axis and a spotcount feature and threshold_30 mask on the x-axis.

**D.** Fraction of polarized cells in the 15 min, 1 hour and 3 hour populations after classification using the three combinations of machine learning classifiers and user-defined features that best resembled the manual quantification (dashed lines). 'Manual' indicates manual quantification after microscopy (data from Fig. 1B) that was used as a template. Only a very small reduction from the 1 hour to the 3 hours time point was observed that did not reflect the strong reduction observed in the template quantification.

### Supplementary Figure S3

**A**

Classification Result

| Truth | Polarized | Unpolarized | Patchy |
|---|---|---|---|
| Polarized | 92.1 | 4.7 | 3.2 |
| Unpolarized | 1.1 | 97.7 | 1.2 |
| Patchy | 5.6 | 1.1 | 93.3 |

**B**

| | **Precision (%)** | **Recall (%)** | **F1 (%)** |
|---|---|---|---|
| Polarized | 97.5 | 92.1 | 94.7 |
| Unpolarized | 94.9 | 97.7 | 96.3 |
| Patchy | 83.3 | 93.3 | 88.0 |
| Total | 94.9 | 94.7 | 94.7 |

**Supplementary Figure S3: Evaluation of the polarized, unpolarized and patchy separation.**
**A:** Confusion matrix generated by Amnis AI depicting the percent of cells from the truth populations (rows) classified as polarized, unpolarized, or patchy (columns). Diagonal, green boxes represent correctly classified cells, white/pink boxes represent cells wrongly classified as the group of that column.
**B:** Table of Precision, Recall and F1 values calculated as described in the methods section for the separation of polarized, unpolarized and patchy cells. Precision determines the accuracy of positive preditions made by the model (taking false positives into account), Recall quantifies the ability of the model to identify all relevant cases (taking false negatives into account), the F1 score is a weight-ed average that considers both false positives and false negatives.

### Supplementary Figure S4

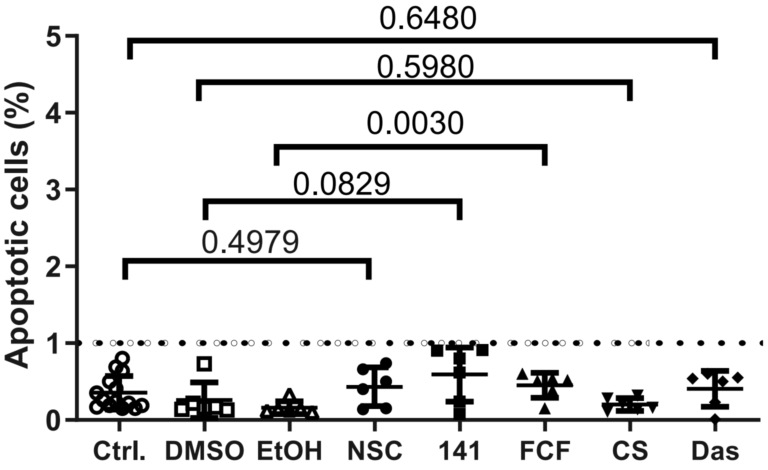


**Supplementary Figure S4: Fraction of apoptotic cells in experiments shown in Figure 3B and 3D.**

As last step in our gating strategy, before classification of cells, apoptotic cells were removed from the IFC analysis (Suppl. Fig. S1F). The amount of apoptotic cells was determined as described by Henery et.al., 2008). The fraction of apoptotic cells for the experiments shown in Fig. **3B** and **3D** are shown as %, the cutoff was set to 1%.

### Supplementary Figure S5

**A**

Classification Result

| Truth | Cytoplasmic | PM-localized |
|---|---|---|
| Cytoplasmic | 95.6 | 4.0 |
| PM-localized | 4.4 | 96.0 |

**B**

| | **Precision (%)** | **Recall (%)** | **F1 (%)** |
|---|---|---|---|
| Cytoplasmic | 96.6 | 99.7 | 98.2 |
| PM-localized | 99.7 | 95.9 | 97.8 |
| Total | 98.0 | 98.0 | 98.0 |

**Supplementary Figure S5: Evaluation of the cytoplasmic and PM-localized separation.**
**A:** Confusion matrix generated by Amnis AI depicting the percent of cells from the truth populations (rows) classified as cytoplasmic or PM-localized (columns). Diagonal, green boxes represent correctly classified cells, pink boxes represent cells wrongly classified as the group of that column.
**B:** Table of Precision, Recall and F1 values calculated as described in the methods section for the separation of cytoplasmic and PM-localized cells. Precision determines the accuracy of positive preditions made by the model (taking false positives into account), Recall quantifies the ability of the model to identify all relevant cases (taking false negatives into account), the F1 score is a weighted average that considers both false positives and false negatives.

#### Supplementary Methods:

##### 1. Gating Strategy:

After compensation, all images were gated in Amnis IDEAS v.6.3. An overview of the features and masks used in the gating strategy is shown in Table 1, an example of the gating strategy can be found as Supporting Information in Supplementary Figure S1.

| Feature | Mask | Feature description | Used for |
|---|---|---|---|
| Gradient RMS_M01_Ch01 | M01 a default mask set to capture all pixels above background in the brightfield images | Determines image sharpness by comparing adjacent pixels | Gating on cells in focus |
| Area_M01 | M01 | Determines the area of the mask | Gating on single cells |
| Aspect Ratio_M01 | M01 | Determines the ratio between height and width the mask | |
| Spot Count_ Morphology(M07, Ch07)_4 | Morphology(M07,Ch07) a default mask set to capture all pixels above background in channel 7 (DAPI) | Counts the number of areas defined by the mask for each cell | Count the number of nuclei |
| Raw Max Pixel_MC_Ch02 | MC a mask set to capture all pixels that have a signal above background in any channel. | Determines the pixel value of the pixel with the highest intensity within the mask. | Gating on cells without over-saturated pixels in channel 2 (GFP) |
| Intensity_MC_Ch02 | MC | Determines the sum of pixel values after back-ground subtraction | Gating on GFP-positive cells |
| Bright Detail Intensity R3_Threshold(M07, Ch07, 30)_Ch07 | Threshold(M07,Ch07,30) covers pixels above background in channel 7 with the 30 % highest intesity | Measures the intensity of localized bright spots within the mask. This was used to detect spotty nuclei. | Excludes apoptotic cells by leaving out cells with small spotty nuclei (Henery et al., 2008) |
| Area_Threshold(M07, Ch07, 30) | Threshold(M07,Ch07,30) | Measures the area of the mask. This was used to find small nuclei. | |

**SupplementaryTable 1: Overview of features and masks used in the gating strategy.**
The table lists the features and masks used in the gating strategy applied before AI analysis, together with a short description of the feature and what it was used for during gating. An example for the gating strategy is shown in Supplementary Figure 1.

The predefined features Gradient RMS, Area and Aspect ratio were used with an M01 mask to identify cell images in focus and then single cells. To remove remaining doublets or double nucleated cells, a morphology mask in Channel 7 was combined with the spot count feature which counts the number of separated mask-areas per cell (Spot Count_Morphology(M07, Ch07)_4) to gate events with one nuclear spot. The Raw Max Pixel feature was used to exclude cells with oversaturated pixels (Raw Max Pixel_MC_Ch02>4094) in channel 2 (GFP), before gating on GFP positive cells (Intensity_MC_Ch02>10,000). The cutoff for this gate was based on a sample with untransfected (GFP negative) cells. To exclude apoptotic cells, a threshold mask in channel 7 covering the area with 30 % highest DAPI intensity was combined with the feature Bright Detail intensity R3 (Bright Detail Intensity R3_Threshold(M07, Ch07, 30)_Ch07)). This feature was plotted against an Area feature using the same threshold mask (Area_Threshold(M07, Ch07, 30)), as described by (Henery et al., 2008), and non-apoptotic cells were gated as the final population. Approximately 50 % of cells in each sample were included for further analysis after gating.

**2. Classification by manual adjustment and machine learning:**

As a template, we used the data generated by the microscopy-based method shown in Fig. 1B. The measurements for the three time points (15 minutes, 1 hour and 3 hours) were pooled to generate 'merged data', which was used for classification by help of IDEAS machine learning and AI modules. Initially, 5 truth populations were generated from merged data consisting of polarised cells with one pole, polarised cells with two poles, unpolarised cells with plasma membrane ezrin localisation, unpolarised cells with cytoplasmic ezrin localisation and "patchy" referring to several small ezrin spots of different intensity. The patchy localisation was introduced as there was a small but consistent fraction of cells showing this phenotype that could not be consistently classified as either polarised or unpolarised by the software. Importantly, the manual, microscopy-based method did not discriminate this additional class. For the final analysis, the two polarized and the two unpolarized classes were combined into one polarized and one unpolarized class.

We first attempted to classify polarised and unpolarised cells using the machine learning module in IDEAS (Supplemental Fig. S2). However, separations based on the machine learning-generated classifiers did not sufficiently reflect the manual quantification method, as it overscored the fraction of polarised cells at the three hours time point, resulting in no further decrease of polarisation from the 1 hour to 3 hours time point (Supplemental Fig. S2B).

In order to improve the analysis, the machine learning-generated classifiers were combined with various user-defined features available in IDEAS (Supplemental Figure 2 C). The combinations that we tested only slightly improved the outcome when applied to the test dataset of 15 minutes, 1 hour and 3 hours time points (Supplemental Fig S2D). Eventually, we employed the correction functions in the AI module, which allowed us to establish the final classification algorithm as described in the main Methods and Results sections (Fig. 1D).

**Supplementary Methods References:**

Henery, S., George, T., Hall, B., Basiji, D., Ortyn, W., & Morrissey, P. (2008). Quantitative image based apoptotic index measurement using multispectral imaging flow cytometry: A comparison with standard photometric methods. *Apoptosis: An International Journal on Programmed Cell Death*, *13*(8), 1054–1063. https://doi.org/10.1007/s10495-008-0227-4
